# GIC: A computational method for predicting the essentiality of long noncoding lncRNAs

**DOI:** 10.1101/177923

**Authors:** Pan Zeng, Ji Chen, Yuan Zhou, Jichun Yang, Qinghua Cui

## Abstract

Measuring the essentiality of genes is critically important in biology and medicine. Some bioinformatic methods have been developed for this issue but none of them can be applied to long noncoding RNAs (lncRNAs), one big class of biological molecules. Here we developed a computational method, GIC (Gene Importance Calculator), which can predict the essentiality of both protein-coding genes and lncRNAs based on RNA sequence information. For identifying the essentiality of protein-coding genes, GIC is competitive with well-established computational scores. More important, GIC showed a high performance for predicting the essentiality of lncRNAs. In an independent mouse lncRNA dataset, GIC achieved an exciting performance (AUC=0.918). In contrast, the traditional computational methods are not applicable to lncRNAs. As a public web server, GIC is freely available at http://www.cuilab.cn/gic/.

## INTRODUCTION

Essential genes constitute a small fraction in a genome of an organism. However, these genes underpin numerous core biological processes and are indispensable for cell viability. Insufficient expression of essential genes will lead to increased vulnerability and loss-of-function mutations of essential genes often cause lethal phenotypes (Korona, 2011; Peters, et al., 2016). Hence the classification of genes as either essential or non-essential for organism survival has a profound influence on the study of molecular basis of various biological process (Wang, et al., 2015), disease genes, drug targets, and genome design (Liu, et al., 2015). In recent years, efficient gene knockout or knockdown by CRISPR/Cas9 and RNAi have been widely used to systematically evaluate the essentiality of genes and lncRNAs(Evers, et al., 2016; Morgens, et al., 2016) in whole organisms (Peters, et al., 2016) and human cells (Blomen, et al., 2015; Wang, et al., 2015; Wang, et al., 2014; Zhou, et al., 2014; Zhu, et al., 2016). These studies provided great helps in identifying functionally important genes and thus have great potential in discovering new genes for disease therapy and diagnosis (Tzelepis, et al., 2016). However, the problem is that these techniques are normally time and labor consuming and hard to be applied to mammals in a large-scale.

Therefore, computational methods have been developed as an effective complement of the experimental approaches. Sequence conservation score measured by comparative genomics and degree in a protein-protein interaction (PPI) network were proposed to evaluate gene essentiality based on the observation that these metrics show significant correlations with gene essentiality (Liang and Li, 2007). In addition, machine learning based method was also developed (Cheng, et al., 2013; Deng, et al., 2011; Seringhaus, et al., 2006). Moreover, more complex topology features of PPI network are also be used for the prediction of essential genes(del Rio, et al., 2009; Gatto, et al., 2015; Li, et al., 2016; Li, et al., 2015; Peng, et al., 2015). More recently, a nucleotide-based computational method was developed for the prediction of human essential genes(Guo, et al., 2017). These methods provided great helps for identifying essential protein-coding genes. Nevertheless, these methods require attributes discriminating essential genes, e.g. gene expression, conservation, sequence features, gene ontology (GO) annotation, protein domain, protein subcellular location, and interaction network topological properties. Therefore, the problem is that these methods often fail to predict the essentiality of lncRNAs, a big class of RNA molecules identified recently in human genome(Iyer, et al., 2015; Zhao, et al., 2016). The reason is that information needed by these methods is usually unavailable for lncRNAs because most lncRNAs show low sequence conservation, low expression level, and high specificity (Iyer, et al., 2015). Moreover, for most of the lncRNAs, information such as the ontology, subcellular location, and interaction network topological properties are still not available (Iyer, et al., 2015). In addition, although Pheg(Guo, et al., 2017) is designed based on RNA sequences but it did not work well on lncRNAs. The reason could be it is based on the CDS sequences of an mRNA.

Here, we developed GIC (Gene Importance Calculator), an algorithm that can efficiently quantify the essentiality of lncRNAs. In addition, besides lncRNAs, GIC also works on protein-coding genes. The results showed that GIC has good performance in predicting essential protein-coding genes. More importantly, GIC showed a high performance in predicting essential lncRNAs, whereas other methods can not be applied to lncRNAs. According to our knowledge, GIC is the first computational method to predict essential lncRNAs. GIC web server and the source code are freely available at http://www.cuilab.cn/gic.

## MATERIALS AND METHODS

### Datasets of RNA sequences

We downloaded human (GRCh37/hg19; Nov 9, 2014) and mouse (GRCm38/mm10; Jan 8, 2015) mRNA sequences deposited in the UCSC Table Browser (Karolchik, et al., 2004). Human and mouse lncRNA transcripts were downloaded from the NONCODE database (Zhao, et al., 2016) (version 4) and the sequences longer than 200 nt were retained.

### Datasets of essential genes

We retrieved human and mouse essential protein coding genes from DEG(Luo, et al., 2014) (version 10). In addition, we collected 7 mouse essential lncRNAs and 7 non-essential lncRNAs with experimental evidence as an independent testing set. These lncRNAs were annotated according to the Mouse Genome Informatics (MGI) database (Bello, et al., 2015; Bult, et al., 2016) (http://www.informatics.jax.org/phenotypes.shtml) and the results from Sauvageau et al.’s assays (Sauvageau, et al., 2013). Gene CRISPR/Cas9 scores in the KBM7 cell line were obtained from *Wang et al.*’ study (Wang, et al., 2015).

### RNA sequence features

The first feature is RNA sequence length. Then, with a sliding window of length 3 and step size of 1, we counted the number of times each of the 64 nucleotide triplets (e.g., ACT, GCC) occurred *c*_*i*_ and converted it to frequency *f*_*i*_ by the following formula.

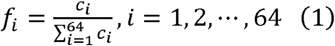

Besides, we used RNAfold (Hofacker, et al., 1994) (version 1.8.5) to predict RNA secondary structure with default parameters and calculated the minimum free energy (MFE) of the secondary structure. Given that longer RNAs favor lower energy state, we introduced here normalized MEF (nMFE) as follows,

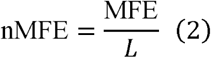

where *L* is RNA sequence length. We then mapped the RNA sequence features to their corresponding genes. For genes with multiple transcripts, we used the mean value in subsequent analysis. The ID mapping files was retrieved from the Ensembl database (Yates, et al., 2016) (release 83) with the R/Bioconductor package biomaRt (Durinck, et al., 2009) and manually curated.

### Logistic regression model and GIC score

To reduce the number of features, especially nucleotide triplet features, we ranked the nucleotide triplet features according to their individual AUC and retained only the top five nucleotide triplet features (CGA, GCG, TCG, ACG, TCA; the same for both human and mouse) without severe co-linearity problem (Pearson correlation < 0.8) with other nucleotide triplet features. Moreover, considering that negative samples greatly outnumbered positive samples in the training set, a subset of negative samples was randomly selected to keep a 1:1 positive-to-negative ratio in the training dataset. Nevertheless, all negative samples were retained in the testing datasets in order to reflect the realistic performance of GIC score. After that, logistic regression models were constructed and cross validated for human and mouse genes and mouse lncRNAs separately. The logistic regression model is that

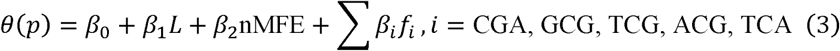

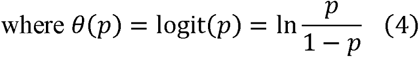

*β*s are the coefficients of corresponding model and *p* is the conditional probability that a gene is essential (*Y* = 1). Accordingly, we defined the GIC score as the probability output *p* of the corresponding logistic regression model. That is

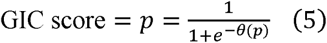

### Correlation analysis between GIC score and well-established measures of essential genes

To explore the relationship between GIC score and several known measures of essential genes, we downloaded corresponding datasets described in detail below and got the intersections of GIC scores and each of them. To assess gene persistence, we counted the homolog number for each gene using data from the Homologene database (NCBI Resource Coordinators., 2016) (build 68). To evaluate sequence conservation, we retrieved the dN/dS ratio of each one-to-one mouse-human (and human-mouse) ortholog pair from the Ensembl database (release 83). The interaction network degrees were derived from the protein-protein interactions recorded in the BioGRID database (Stark, et al., 2006) (release 3.4.135). At last, genes were sorted by GIC score and median-binned into 200 bins for clearer illustration.

### Comparing the accuracy of human and mouse essential gene prediction

Gene essentiality was annotated as a Boolean value based on the corresponding essential gene set acquired from DEG. Using the R package pROC (Robin, et al., 2011), the ROC curves were plotted and the AUC values for GIC score and the abovementioned measures were calculated and compared. Note that only the samples for which all of the above-mentioned measures were available were used during the comparison.

### Cell culture

Human T/G HA VSMC Cells were cultured in DMEM medium supplemented with 15% FBS, 2mM L-glutamine, 100 U/mL penicillin, and 10 mg/mL streptomycin. The media were renewed twice a week. All experimental procedures were conducted within a CO_2_ incubator at a temperature of 37°C, in an atmosphere of 95% air and 5% CO_2_.

### siRNA knockdown of target mRNAs in T/G HA VSMCs

T/G HA VSMCs (vascular smooth muscle cells) with the confluence of 60% were synchronized with serum-free starvation for 24 hours, and then transfected with siRNA mixtures against various mRNAs (50nM) or scrambled siRNA (50nM) using VigoFect transfection kit (Vigorous Biotechnolog, Cat No T001) for 48 hours. The siRNA mixture was transfected into T/G HA VSMC cells according to the manufacturer’s siRNA gene silencing protocol. Basically, the siRNAs against each target mRNA were the mixture of four sets of sequences according to different part of target mRNA. All the siRNA sequences were designed and synthesized by Beijing Biolino Inc. All the siRNA sequences against various target mRNAs were provided in **Supplementary Table S1**. The scrambled siRNA was also provided by Beijing Biolino Inc.

### Real time PCR analysis of target mRNAs after siRNA transfection

T/G HA VSMCs were transfected with 50nM siRNAs or scrambled siRNAs as detailed above. Forty eight hours post transfection, total cellular RNA was extracted using the Trizol reagent according to the manufacturer’s instructions. 0.5-1.0μg of total RNA was used for the reverse transcription reaction. Quantitative real time PCR was performed using the DNA Engine with Chromo 4 Detector (MJ Research,Waltham, MA). The relative expression of target genes in various groups were calculated using 2^-ΔΔCt^ methodology. β-actin mRNA was used as housekeeping gene in the current study. All primer sequences used for real-time PCR assays were listed in **Supplementary Table S2**.

### Cell viability assay

Cell viability was measured by MTT assay as detailed previously. In brief, T/G HA VSMCs were seeded and transfected in 24-well plates. At 48 hours post siRNA transfection, MTT was added into the culture medium to the final concentration of 0.5mg/ml, and then the cells were incubated for 4 hour at 37°C in incubator. The culture medium was removed, and cells were lysed by gently rotating in 250uL DMSO for 30 minutes in darkness at room temperature. The absorbance at 490nm was measured using an automatic plate reader. In each experiment, 3-4 observations were set and determined for each siRNA mixture. The average absorbance reflected cell viability with the data normalized to the control group.

### Flow cytometry analysis of cell apoptosis

At 48 hours post transfection, T/G VSMC cells were managed using Apoptosis Detection Kit with 7-AAD according to the manufacturer’s protocol (Biolegend). In brief, the cells were washed with ice-cold PBS twice, and then resuspended in 100μl of Annexin X binding buffer (10mM HEPES, pH 7.4, 140mM NaCl, 1mM MgCl2, 5mM KCl, and2.5 mM CaCl2), and then added 1μl FITC Annex V and 1μl 7-Add viability staining solution. The cells were incubated at room temperature in darkness for 15 minutes, and then were analyzed by FACScan analysis with Cellquest software (Becton Dickinson).

### Code availability

GIC is implemented in Python and it relies on the external program RNAfold. We provide convenient online service on our GIC web server (http://www.cuilab.cn/gic). However, as for large RNAs or batch jobs, we recommend users download the source code on this server. Besides, the pre-calculated GIC scores of human and mouse genes, including both mRNAs and lncRNAs, are also available on the server.

## RESULTS AND DISCUSSION

### The construction of GIC

In brief, we managed to construct a logistic regression model (GIC) by integrating several features that can be derived from RNA sequences or predicted RNA secondary structures for measuring gene essentiality. First of all, the length of a RNA sequence was considered as a feature of gene essentiality based on the observation that RNAs encode conserved proteins are longer than those encode proteins with less conservation(Lipman, et al., 2002). And then we integrated the frequencies of some specific nucleotide triplets into the model. In addition, we found mRNA products of essential genes often form more stable structures, which are found to influence gene expression (Wan, et al., 2014). Thus, we utilized RNAfold (Hofacker, et al., 1994) to predict RNA secondary structure and its minimum free energy (MFE). Given that longer RNAs normally have lower MFE than shorter RNAs, we normalized MFE by sequence length in the model. Finally, given the serious imbalance between the numbers of essential genes and non-essential genes, we randomly selected a subset of negative samples (non-essential genes) to keep a balanced positive-to-negative ratio in the training dataset and trained the logistic regression model based on the balanced dataset. GIC score was defined as the probability output of the model.

### Comparison of GIC method with previous computational methods and experimental methods on protein-coding genes

We first tested GIC score on protein-coding genes. Because we aim to develop a method for predicting essential lncRNAs, we did not compare GIC with each existing methods. For simplicity, we only tested GIC score with three well established scores, protein network degree, dn/ds score, and homolog number score. We observed significant correlations between human GIC scores and other computational scores (Figure 1a-c; Spearman ρ = 0.67, P = 6.17×10^-27^ with homolog number, Spearman ρ = –0.92, P = 0 with dN/dS, Spearman ρ= 0.69, P = 4.51×10^-30^ with protein interaction network degree, respectively). For mouse genes, we got similar results (Figure 2a-c).

**Figure 1.**
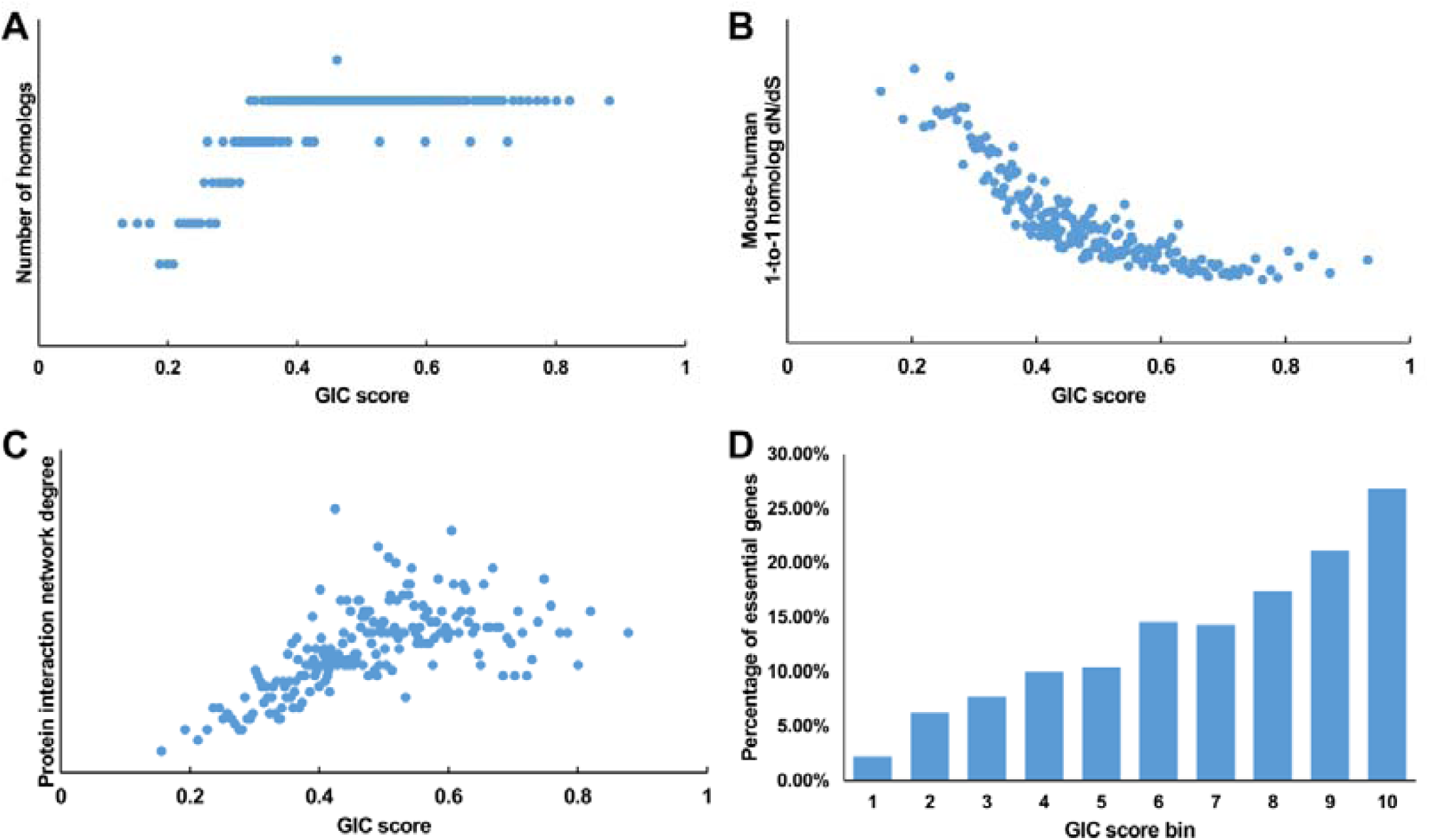
Correlation of GIC score with known measurements of essentiality in human genes. (a) Genes with higher GIC scores tend to have more homologs across species. (b) Genes with higher GIC scores tend to have slower evolutionary rate as measured by dN/dS ratio. (c) Proteins encoded by genes with higher GIC scores tend to have higher degrees in protein interaction network. (d) The percentage of human essential genes increases with GIC score.

**Figure 2.**
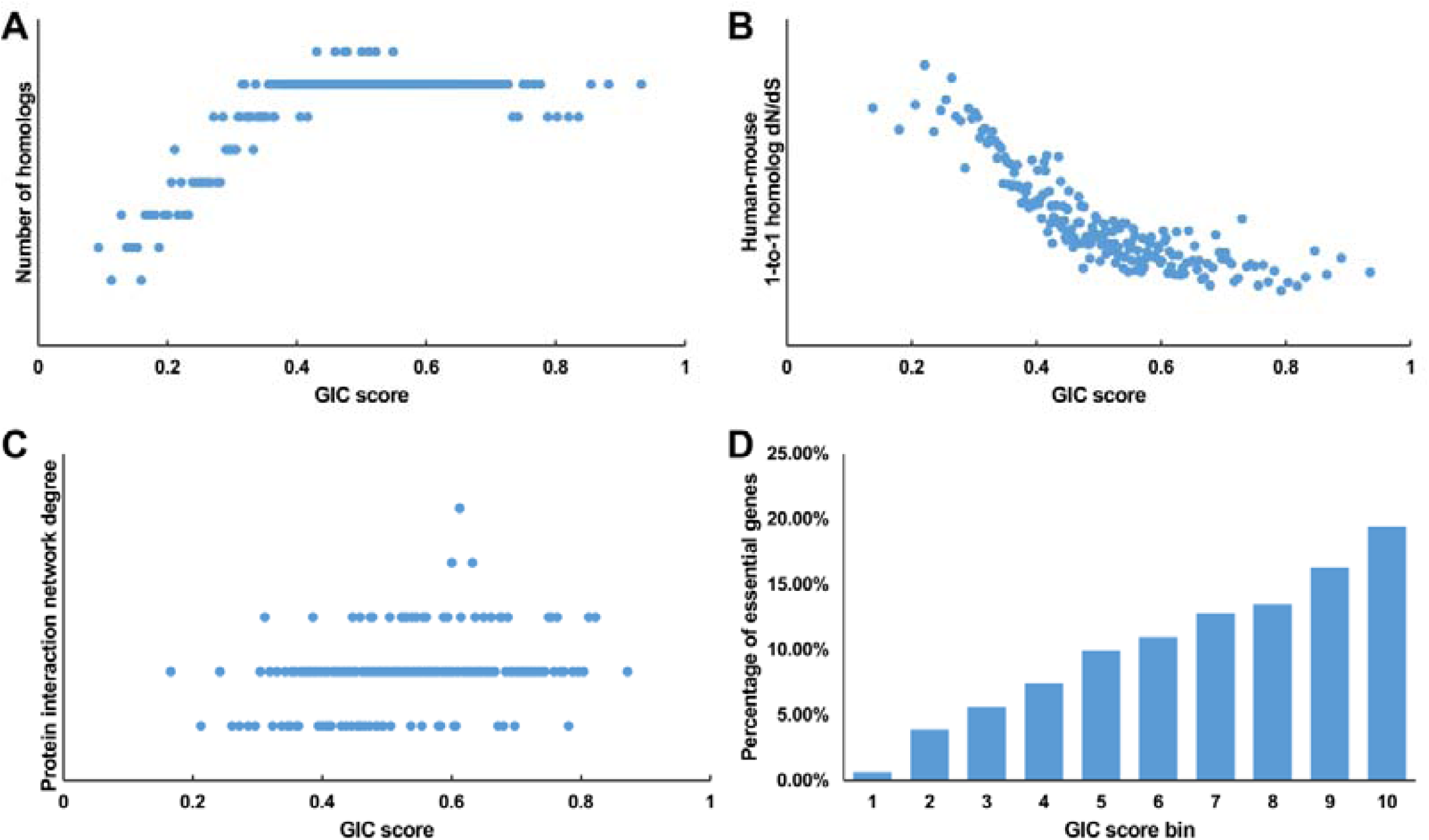
Correlation of GIC score with known measurements of essentiality in mouse genes. (a) Genes with higher GIC scores tend to have more homologs across species. (b) Genes with higher GIC scores tend to have slower evolutionary rate as measured by dN/dS ratio. (c) Proteins encoded by genes with higher GIC scores tend to have higher degrees in protein interaction network. (d) The percentage of human essential genes increases with GIC score.

Furthermore, we took the human and mouse essential genes stored in the DEG database as the benchmarks to evaluate the accuracy of GIC score. First, we ranked the human and mouse genes by GIC score and simply divided them into ten equal groups, respectively. Indeed, essential genes were enriched in groups of genes with higher GIC scores for both human (Figure 1d; P = 1.31×10^-69^, Pearson’s Chi-squared test) and mouse (Figure 2d; P = 7.80×10^-68^, Pearson’s Chi-squared test).

In addition, in recent years, genetics-based methods such as CRISPR/Cas9 have been used for identifying essential genes in a specific condition, for example some cancer cell line. To test whether CRISPR/Cas9 scores in one condition can be useful in other conditions, we also compared GIC score with CRISPR/Cas9 scores. As a result, in terms of performance on the area under the receiver operator characteristic (ROC) curve (AUC), GIC score has an AUC of 0.675), whereas CRISPR/Cas9 scores in KBM7 cell line (Wang, et al., 2015) has an AUC of 0.568 (Figure 3a), suggesting that CRISPR/Cas9 scores in one condition could be not useful for evaluating the essentiality of genes in other conditions. Moreover, GIC score showed competitive performance compared with the three well-established computational scores including homolog number (AUC = 0.622, P = 2.76×10^-11^, bootstrap test), the dN/dS score of mouse-human 1-to-1 homolog (AUC = 0.639, P = 2.57×10^-5^, bootstrap test) and protein interaction network degree (AUC = 0.647, P = 1.14×10^-3^, bootstrap test).

**Figure 3.**
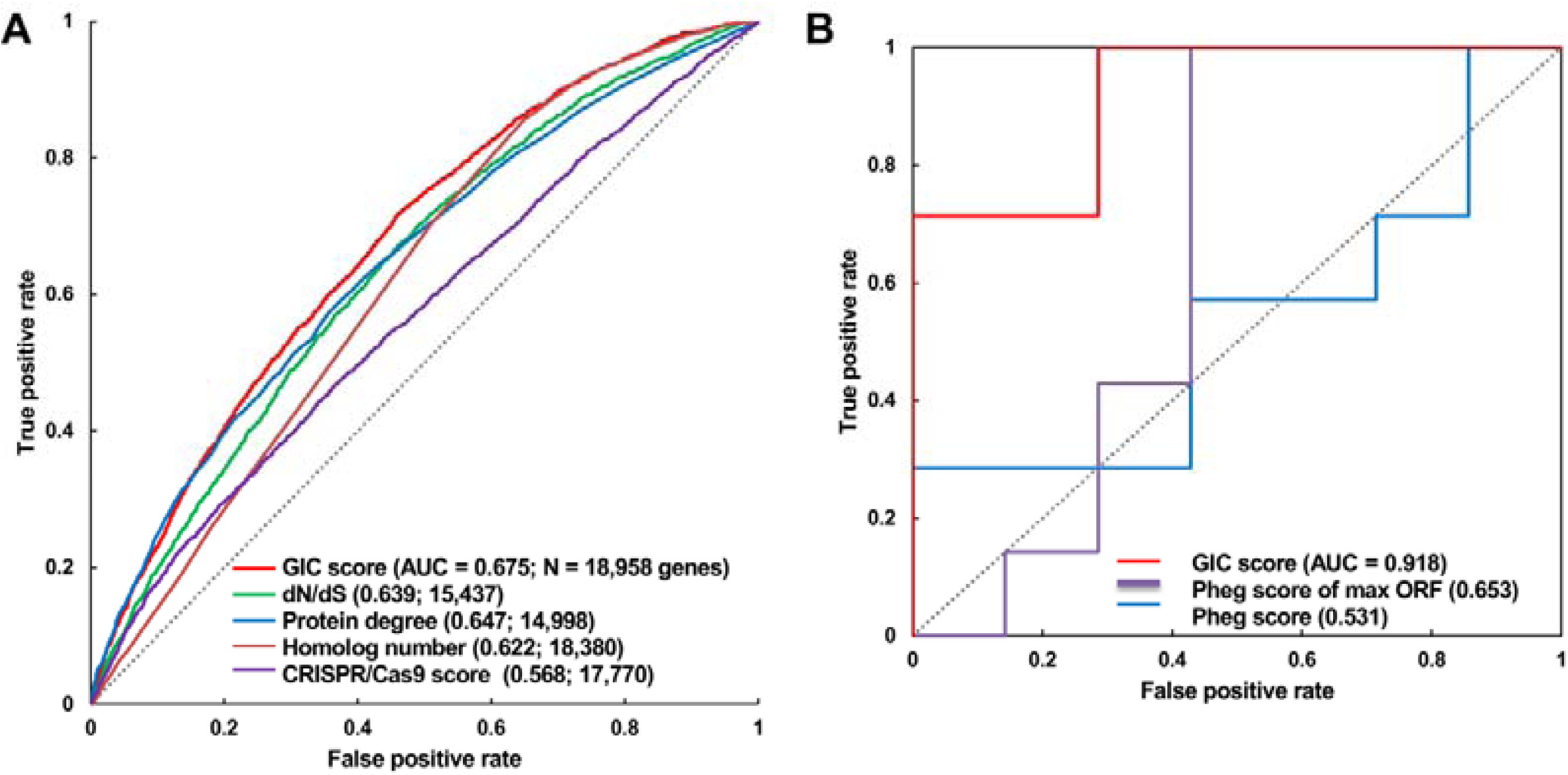
Validation of GIC score. (**a**) ROC curves illustrating the results from human essential gene prediction analysis. (**b**) ROC curves illustrating the results of essentiality prediction in an independent mouse lncRNA dataset.

### Performance of GIC method on predicting the essentiality of lncRNAs

Next, we directly tested if GIC score is feasible to predict essential lncRNAs. To this end, we gleaned 14 mouse lncRNAs, of which seven were essential and the others were non-essential in mutagenesis assays, as an independent testing dataset (**Methods**). On this testing lncRNA dataset, GIC score showcased an AUC of 0.918 (Figure 3b and **Supplementary Table S3**). The outcome again verified the viability of GIC score. Currently, there is no specific tool for essential lncRNA prediction, mainly due to the special characteristics of lncRNAs. Our GIC score can measure lncRNA essentiality with RNA sequence only and will serve as a promising tool to prioritize functionally important lncRNAs. Given that Pheg also only needs sequences as the input data, here we compared GIC with Pheg. For each lncRNA, we first inputed its whole lncRNA sequence. Because Pheg works on mRNA CDS regions, we run Pheg using the putative maximum CDS sequence, which was predicted using ORF Finder (https://www.ncbi.nlm.nih.gov/orffinder/). As a result, Pheg did not achieve a comparable performance for both whole sequences (AUC =0.531) and CDS sequences (AUC =0.653, Figure 3b).

### A case study of using GIC to select genes essential in cancer but non-essential in other cells

Genome-level gene knockout in cancer cell lines provides great helps in identifying essential genes in given cells (Wang, et al., 2015). These essential genes could be candidate targets for cancer therapy. For specificity of therapy, one ideal condition is that these essential genes are non-essential in other types of normal cells. This makes it important to identify these genes that are essential in given cancer cells but non-essential in other cells, and also those non-essential in cancer genes and essential in other cells. To test GIC for such studies, we randomly selected 10 genes, which include 5 GIC-predicted essential (among the top 25% GIC scores) but CRISPR/Cas9 non-essential genes in the KBM7 cell line (among the bottom 25% CRISPR/Cas9 scores) (*RIMKLA*, *ATF7IP*, *PFN3*, *HIST1H3J* and *DDIT4*, these genes were named as GIC_essential_-Cas9_nonessential_ genes here) and 5 GIC-predicted non-essential (among the bottom 25% GIC scores) but CRISPR/Cas9 essential genes (among the top 25% CRISPR/Cas9 scores) (*CBWD3*, *SLC35B1*, *CCDC33*, *DPH6* and *ZMAT2*, these genes were named as GIC_nonessential_-Cas9_essential_ genes here). Because blood vessel is a basic tissue for cancer cell survival, we evaluate the effects of the ten genes on VSMCs, which one class of key cells for blood vessel. The efficacy of siRNAs treatment on target mRNA levels were first determined by real time PCR assays, revealing that siRNAs treatment significantly reduced the target mRNA levels by about 60-90% (Figure 4a). For evaluating the essentiality of the 10 target mRNAs on cell survival, they were randomly divided into two groups for siRNAs transfection to reduce experimental errors. It is also noteworthy that the person who performed the experimental validation was blind to the essentiality of these genes during the experiments. MTT assays indicated silencing GIC_essential_-Cas9_nonessential_ genes *RIMKLA*, *ATF7IP*, *PFN3*, *HIST1H3J* and *DDIT4* significantly reduced cell viability by 30-50% (Figure 4b-c). In contrast, among the GIC_nonessential_-Cas9_essential_ genes, silencing of *DPH6* reduced cell viability by only about 10%, whereas silencing of *CBWD3*, *SLC35B1*, *CCDC33* and *ZMAT2* had no significant effect on cell viability as evaluated by MTT assays (Figure 4b-c). siRNAs against *HIST1H3J* and *SLC35B1* mRNAs exhibited the highest repression efficacy among all siRNAs. Silencing GIC_essential_-Cas9_nonessential_ gene *HIST1H3J* significantly reduced cell viability, whereas silencing GIC_nonessential_-Cas9_essential_ gene *SLC353H* had no significant effect on cell viability of human T/G HA VSMCs. siRNAs against other eight target mRNAs exhibited comparable silencing efficacy (Figure 4a). Overall, these results strongly indicated that the significant reduction in cell viability after silencing GIC_essential_-Cas9_nonessential_ genes is not due to repression efficacy of siRNAs (Figure 4a-c).

**Figure 4.**
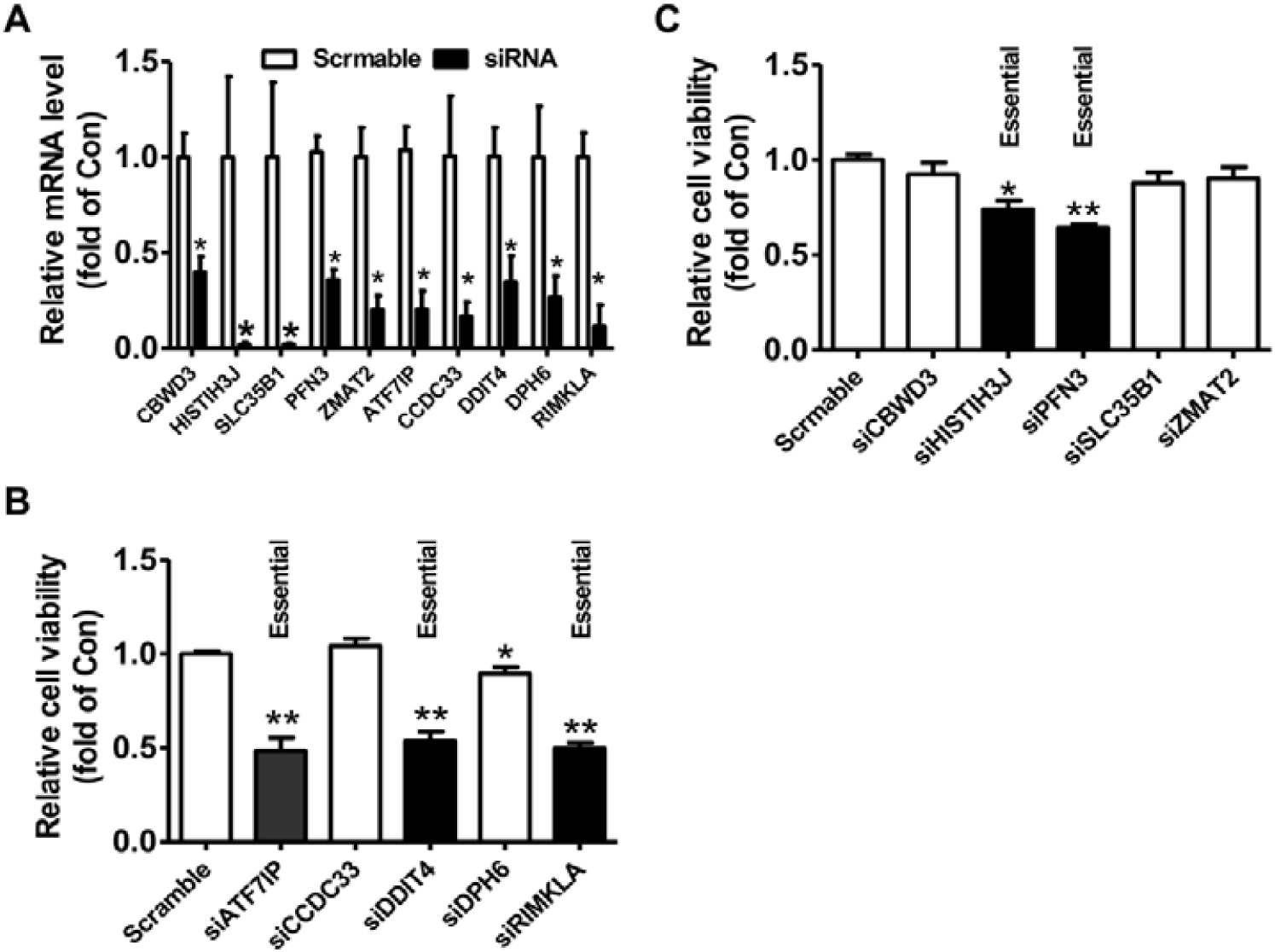
siRNA-mediated silencing of target mRNAs on the cell viability of human T/G HA VSMCs. **(a)** The efficacy of siRNA treatment on the repression of target mRNA levels. The target mRNA levels were analyzed by real time PCR assays at 48 hours post siRNA transfection. N=6, *P<0.05 versus control cells transfected with scrambled siRNAs. **(b-c)** Silencing of target mRNAs on the cell viability. At 48 hours post siRNA transfection, cell viability was determined using MTT assay as described in experimental procedure. In every experiment, 3-4 parallel observations were set for each siRNA mixture. In panels b and c, GIC-predicted essential genes but CRISPR/Cas9 non-essential genes were presented as fill bars, whereas GIC-predicted non-essential genes but CRISPR/Cas9 essential genes were presented as blank bars. N=4, *P<0.05,**P<0.01 versus control cells transfected with scrambled siRNAs.

To further validate the essentiality of GIC_essential_-Cas9_nonessential_ and GIC_nonessential_-Cas9_essential_ genes on cell survival, flow cytometry was performed to analyze the impacts of silencing of two GIC_essential_-Cas9_nonessential_ genes, *RIMKLA* and *ATF7IP*, and two GICn_onessential_-Cas9_essential_ genes, *CBWD3* and *CCDC33*, on the apoptosis of human T/G HA VSMCs. Silencing RIMKLA and ATF7IP markedly increased the proportions of apoptotic cells (51.48±9.38% versus 8.08±1.23% (Control) for *RIMKLA*, and 40.72±7.37% versus 8.08±1.23% (Control) for *ATF7IP*, respectively, P<0.01) (Figure 5a-b). In contrast, silencing *CBWD3* slightly increased apoptotic cells (14.67±1.02% versus 8.08±1.23%, P<0.05), and silencing *CCDC33* had no statistically significant effect on cell apoptosis when compared with control cells (13.43±3.12% versus 8.08±1.23%, P>0.05) (Figure 5a-b). Importantly, there is also significant difference between GIC_essential_-Cas9_nonessential_ and GIC_nonessential_-Cas9_essential_ genes (Figure 5b), confirming the results of the above cell viability assays.

**Figure 5.**
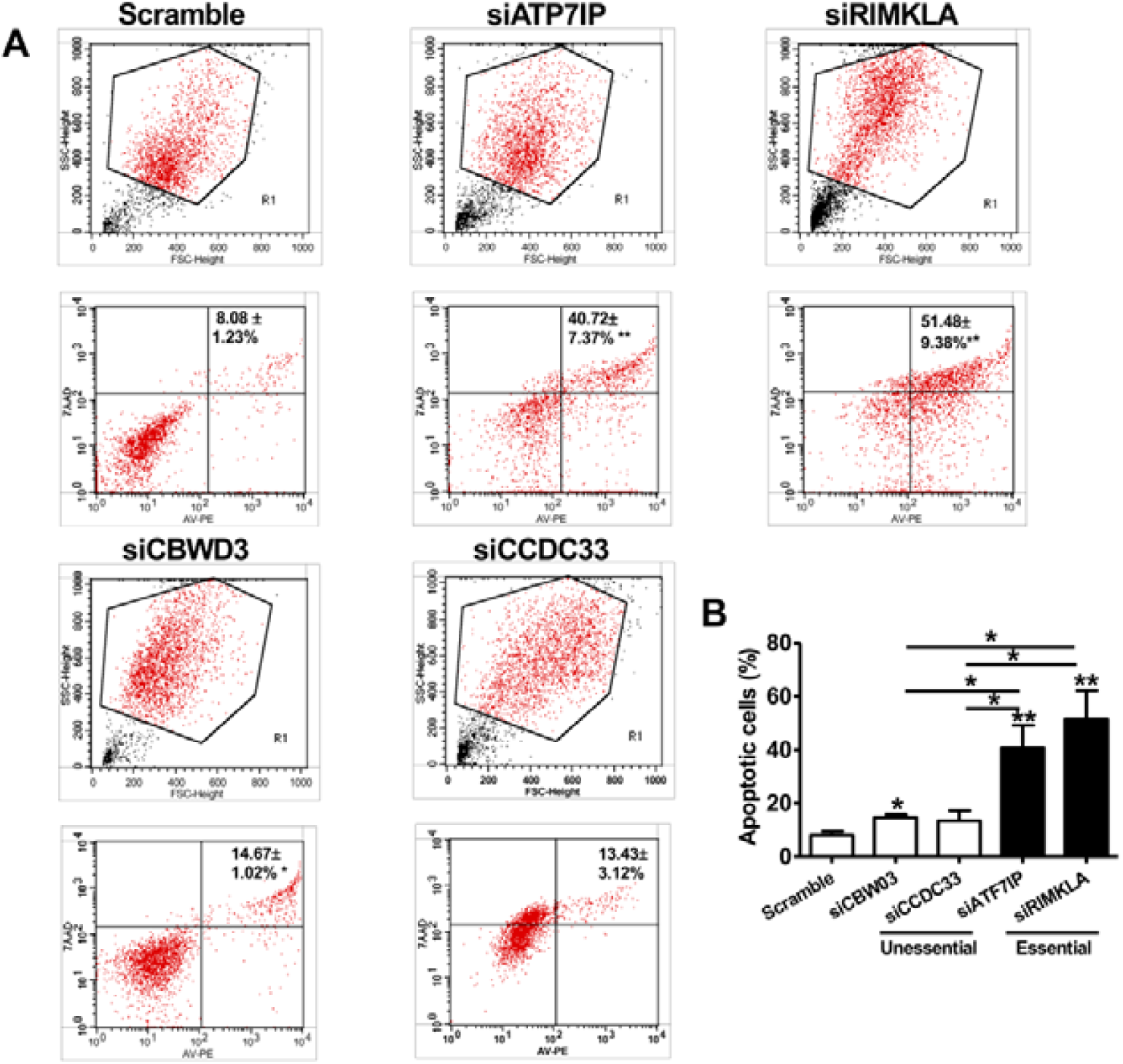
Flow cytometry analysis of apoptosis of human T/G HA VSMCs. The essentiality of two GIC-predicted essential genes but CRISPR/Cas9 non-essential genes and two GIC-predicted non-essential genes but CRISPR/Cas9 essential genes in cell survival were selected for further validation, respectively. The cells were transfected with siRNAs against target mRNAs or scrambled siRNA, and the apoptosis was determined by flow cytometry analysis at 48 hours later. **(a)** Representative images of flow cytometry analysis. **(b)** Quantitative data of apoptosis determined by flow cytometry. N=4, *P<0.05, **P<0.01 versus control cells treated with scrambled siRNAs or between two indicated groups.

As a comparison, Pheg predicted one of the 5 GIC_essential_-Cas9_nonessential_ genes as essential genes and three of the GIC_nonessential_-Cas9_essential_ genes as non-essential genes (**Supplementary Table S4**).

## Discussion

Predicting gene essentiality is an important issue in bioinformatics but computational methods for predicting the essentiality of lncRNAs are still not available. For doing so, here we defined GIC (Gene Importance Calculator) score on the basis of sequence information. Overall, our data validated the high accuracy of GIC score for predicting essential and non-essential genes and lncRNAs. Given that the functions of many human protein-coding genes and lncRNAs are still awaiting exploration, our new method provides an effective strategy for identifying and characterizing new genes and lncRNAs with important functions, which definitely will shed light on the pathogenesis, diagnosis, and therapy of human diseases.

microRNAs (miRNAs) are one class of important small noncoding RNAs. We also tried to test GIC on miRNAs. When evaluating GIC score on miRNAs base on two types of previously presented scores, miRNA conservation score (Wang, et al., 2010) and miRNA disease spectrum width (DSW) score(Qiu, et al., 2012), we found that GIC did not work any more, suggesting the GIC method is not feasible for miRNAs. The reason could be miRNA sequences are very small, so the nucleotide features derived from protein-coding genes are not feasible for miRNAs any more. Finally, because miRNAs also have many species (especially human) specific miRNAs and the number of miRNAs with DSW score is still small, the above two scores for miRNAs are not feasible for all miRNAs. So, it is also important to develop new computational methods to predict essential miRNAs.

## ACKNOWLEDGEMENTS

We thank members of Prof. Cui’s lab and Prof. Yang’s lab for helpful discussions.

